# *De novo* diploid genome assembly for genome-wide structural variant detection

**DOI:** 10.1101/552430

**Authors:** Lu Zhang, Xin Zhou, Ziming Weng, Arend Sidow

## Abstract

Structural variants (SVs) in a personal genome are important but, for all practical purposes, impossible to detect comprehensively by standard short-fragment sequencing. *De novo* assembly, traditionally used to generate reference genomes, offers an alternative means for variant detection and phasing but has not been applied broadly to human genomes because of fundamental limitations of short-fragment approaches and high cost of long-read technologies. We here show that 10x linked-read sequencing, which has been applied to assemble human diploid genomes into high quality contigs, supports accurate SV detection. We examined variants in six *de novo* 10x assemblies with diverse experimental parameters from two commonly used human cell lines, NA12878 and NA24385. The assemblies are effective in detecting mid-size SVs, which were discovered by simple pairwise alignment of the assemblies’ contigs to the reference (hg38). Our study also shows that the accuracy of SV breakpoint at base-pair level is high, with a majority (80% for deletion and 70% for insertion) of SVs having precisely correct sizes and breakpoints (<2bp difference). Finally, setting the ancestral state of SV loci by comparing to ape orthologs allows inference of the actual molecular mechanism (insertion or deletion) causing the mutation, which in about half of cases is opposite to that of the reference-based call. Interestingly, we uncover 214 SVs that may have been maintained as polymorphisms in the human lineage since before our divergence from chimp. Overall, we show that *de novo* assembly of 10x linked-read data can achieve cost-effective SV detection for personal genomes.

## Introduction

Cost-effective whole-genome sequencing has been revolutionized over the past decade by short-read approaches, the most widespread of which have been the consistently improving generations of the original Solexa technology ^1, 2^, now Illumina sequencing. Standard Illumina data supports high-quality, read-mapping-based detection of SNVs (single nucleotide variants) in about 90% of the human genome ^3, 4, 5, 6, 7^. *De novo* assembly on Illumina data has been recognized to be an alternative way to generate comparable SNV and better small indel (insertions and deletions) calls ^8^. However, detection of structural variants (SVs) on the basis of short-fragment Illumina data alone continues to be challenging ^9, 10, 11^, and *de novo* assembly of anything but the simplest microbial genomes ^12^ does not yet generate usefully contiguous genome sequences unless Illumina data is supplemented with other data ^13, 14, 15^.

The lack of long-range contiguity in standard Illumina data has distinct consequences depending on the applications. For SV discovery, split reads and other mapping-based approaches can detect breakpoints but connecting them to call a specific SV remains extremely challenging ^16, 17, 18, 19^. For haplotyping, variants can be phased by population-based methods ^20, 21^ or family-based recombination inference ^22, 23^, but such approaches are only feasible for common SNVs or large pedigrees. Finally, highly polymorphic regions such as the HLA in which the reference sequence does not adequately capture the diversity present in the population are refractory to mapping-based approaches and require *de novo* assembly to reconstruct ^24^; but for *de novo* assembly, short-fragment data are challenged by interspersed repetitive sequences from mobile elements and by segmental duplications, and only supports highly fragmented genome reconstruction ^25, 26^.

In principle, many of the challenges of short-fragment approaches for comprehensive variant discovery can be overcome by long-fragment/read sequencing ^27, 28^. Direct sequencing of long DNA fragments requires single-molecule approaches, such as Pacific Biosciences (PacBio) or Oxford Nanopore (ONT) ^29, 30^, because no enzymatic technology exists that can reliably amplify long DNA fragments of arbitrary sequences. The main trade-offs between Illumina and single-molecule long read approaches can at present be best characterized as low-cost, high base quality, short fragments (Illumina) vs. higher cost, low raw base quality, long fragments (PacBio and ONT)^9, 31^. As a consequence, whole-genome sequencing technologies now tend to be deployed in highly specialized ways that emphasize different methodologies depending on the goal to be achieved: standard 30x Illumina sequencing for small variant detection and relatively low-power SV detection ^7, 32^; mate-pair libraries or single-molecule approaches (i.e., long-fragment) for better SV detection and haplotyping ^9, 33^, and hybrid approaches with many different technologies for *de novo* assembly ^15, 34^.

10X linked-read sequencing provides an alternative way to integrate the advances of short-read and long-fragment sequencning. Novel computational approaches leveraging the special characteristics of 10x Genomics data have already generated significant advances in power and accuracy of haplotyping ^35, 36^, cancer genome reconstruction ^37, 38^, metagenome assemblies ^39^, and *de novo* assembly of human and other genomes ^14, 40, 41^, compared to standard Illumina short-fragment sequencing. 10x linked-read sequencing combines low per-base error and good small-variant discovery with long-range information for much improved SV detection in mapping-based approaches ^38, 42^, and the possibility of long-range contiguity in *de novo* assembly ^40, 41, 43^.

We therefore assessed the ability of *de novo* 10x assemblies on SV detection. Our analyses are based on pairwise alignment of the assemblies’ contigs to the reference genome and finding gaps, a procedure whose compelling simplicity is only possible with assembly-based approaches ^8^. We use a variety of metrics (SVs shared between individuals, support by PacBio data, and alignment to Ape genomes) to assess the accuracy of our assembly-based SV calls. Additionally, we explore the difference between the SV calls and the molecular mechanism that produced the derived allele and are able to identify the true molecular event that brought about a subset of SVs. Finally, we uncover an unexpected number of SVs that have most likely been maintained as polymorphisms since before the last common ancestor of chimps and humans.

## Results

### Library preparation, physical parameters, and sequencing coverage

We prepared and sequenced six whole-genome libraries with diverse total input DNA and fragment size distributions, three for NA12878 and NA24385 each (**Methods**). Accordingly, the data varied in physical fragment coverage (*C*_*F*_), read coverage per fragment (*C*_*R*_), and average fragment size (*μ*_*FL*_) (**Table S1**). Each library was sequenced by three lanes of short-reads to guarantee a sufficient sequencing coverage for assembly. We used SuperNova2 ^40^ for assembly, limiting the sequencing coverage by subsampling to include 1200M reads, or approximately 56X (R_1_ to R_6_ in **Table 1**). The contigs from the six assemblies were aligned against the human reference genome (hg38) to identify SNVs, and indels of 50bp or greater (**Methods**).

**Table 1.** Summary of the assemblies of the six libraries from NA12878 and NA24385. Contigs are aligned to human reference genome (hg38) to calculate the overall genomic coverage and the genomic regions in diploid and haploid states.

| Library | Sample | Contig<br>N50/NA50 (kb) | Scaffold<br>N50/NA50 (Mb) | Coverage<br>(%) | Diploid<br>Regions (%) | Haploid<br>Regions (%) |
| --- | --- | --- | --- | --- | --- | --- |
| $R_1$ | NA12878 | 141.2/116.8 | 27.86/13.43 | 91.9 | 58.9 | 27.7 |
| $R_2$ | NA12878 | 114.9/100.4 | 17.22/6.96 | 91.1 | 73.3 | 11.3 |
| $R_3$ | NA12878 | 99.4/86.3 | 7.93/4.77 | 91.7 | 77.2 | 9.2 |
| $R_4$ | NA24385 | 101.2/89.2 | 8.76/4.66 | 91.3 | 73.4 | 12.2 |
| $R_5$ | NA24385 | 58.4/54.2 | 2.85/1.94 | 91.7 | 79.2 | 5.8 |
| $R_6$ | NA24385 | 129.2/110.3 | 48.66/12.57 | 91.7 | 78.1 | 7.9 |

### Concordance and accuracy of assembly-based SNV calls

We first analyzed SNV calls from the pairwise alignments in order to assess the overall feasibility of assembly-based variant calling. The number of SNV calls from five libraries (R_2_ to R_5_) were comparable, around three million (**Table S2**). By contrast, R_1_ covered the lowest percentage of diploid regions (58.9%) and generated the smallest SNV set (2,635,173, **Table 1** and **Table S2**). The assemblies of each library from the same individual shared more than 92% of SNVs with another, and 2 to 2.4 million SNVs were shared by all the three (**Table S3** and **Figure S1**). Genotype concordances were high for those SNVs shared by all three assemblies of the same individual, more than 99.9% (**Table S4**). These assembly-based calls cover 92.4%-93.6% (NA12878) and 95.1%-96.5% (NA24385) of SNVs called by barcode-aware, mapping-based calls. Genotype concordance between assembly- and mapping-based calls was high for all the libraries, around 99.8% (**Table S5**). Furthermore, we compared assembly-based calls with the ‘gold standard’ Genome In A Bottle (GIAB) call set ^44^. We only evaluated the ‘gold standard’ SNVs that fell within the overlap of diploid regions of our assemblies and of high confidence regions from GIAB (**Methods**). Around 93%-97% of these SNVs (**Table S6**) could be detected by assembly-based calls (**Figure S2**).

We also investigated whether the parameters of library preparation and sequencing might explain some of the differences in SNVs detection among different libraries (**Table S1**). For NA12878 and NA24385, the two libraries with the lowest physical coverages of *R*_2_ and *R*_5_ (*C*_*F*_ =123X and 208X), leading to the worst performance on SNV calling (highest false negative rates and lowest genotype concordance), which was improved if *C*_*F*_ was increased substantially (**Table S6**). We did not observe much difference between *R*_4_ and *R*_6_, suggesting the performance of SNV calls would not dramatically change if the physical coverage was sufficiently high (*C*_*F*_ =803X). The most common assembly-based genotyping errors were heterozygous SNVs miscalled as homozygosity (**Table S6**).

### SV calls from diploid contigs

We inferred large and mid-size indels (>=50bp) from the same contig-to-reference alignments that were used for SNV calling (**Methods**). Two to three times more deletions than insertions were detected in the six assemblies (**Table S2**). The size distributions of different libraries were comparable and a peak was observed near 300bp, in which most of the structural variants were further recognized as Alu sequences (**Methods**, **Figure 1** and **Figure S3**-**S4**). We also observed peaks around 6kb in deletions, where some of the structural variants were annotated as long interspersed nuclear elements (L1s, **Methods, Figure 1** and **Figure S3**-**S4**). SV calls in the three assemblies from the same individual differ somewhat with each assembly having around 30%-40% unique calls, and overlapping calls also constitute similar proportion for each library (**Figure S5).**

**Figure 1.**
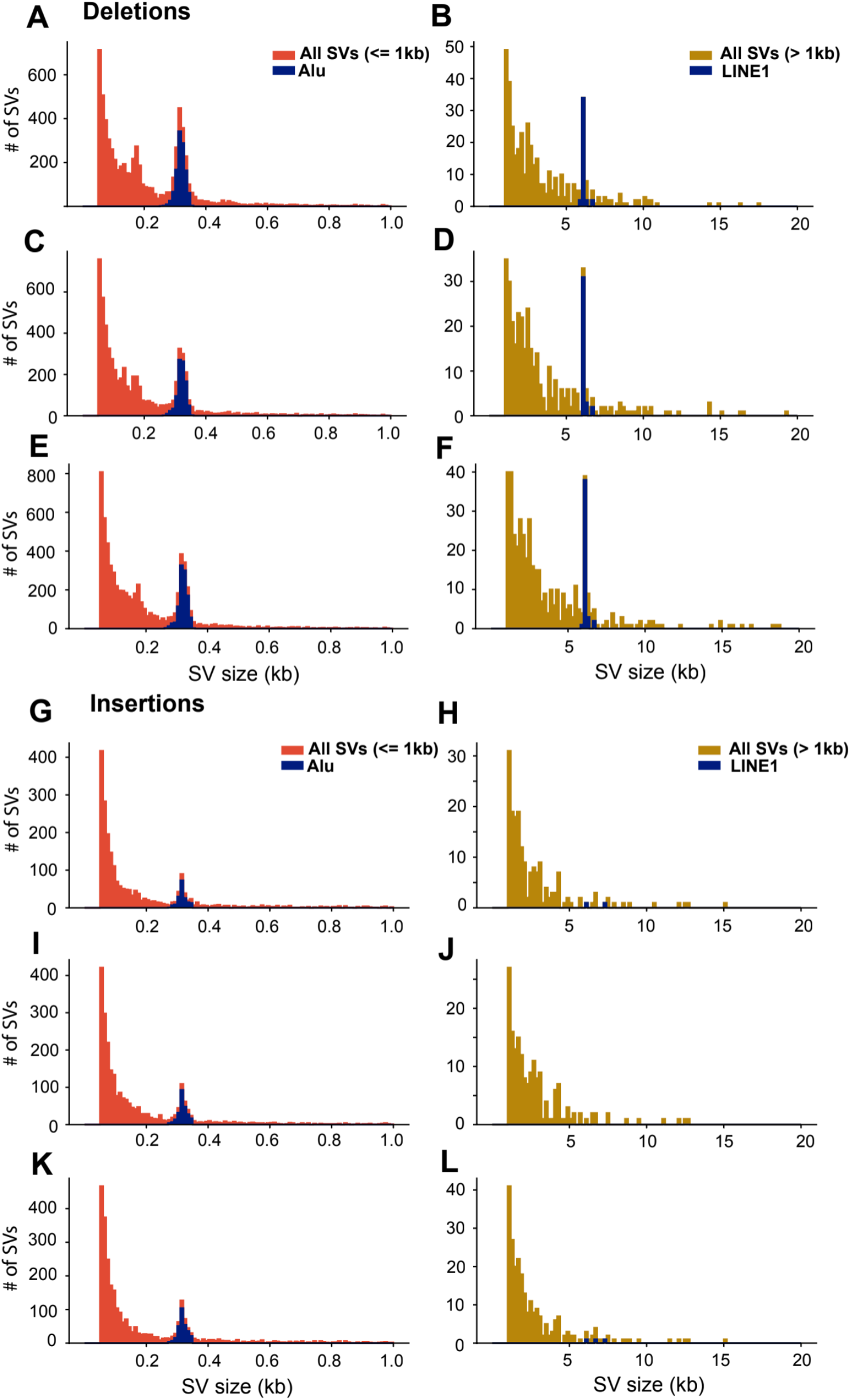
Deletion and Insertion size distributions of NA12878 for *R*_1_ (A, B, G and H), *R*_2_ (C, D, I and G) and *R*_3_ (E, F, K and I).

### Validation of SV calls

Because there is currently no widely accepted ground truth of SVs for the two individuals (NA12878 and NA24385), we designed three criteria to evaluate our calls: supporting evidence from PacBio reads analyzed by svviz2 ^45^; overlap between the two individuals; and finally, by alignment to two ape genomes (chimp and orang; **Methods**; **Figure S6**). For validation analysis, we pooled all non-redundant calls from the three libraries for each individual. This inflates the false positive rate but allows for a more comprehensible analysis. By using the union of the abovementioned three criteria, we could validate roughly half of the deletions (51.3% for NA12878 and 50.7% for NA24385) and almost 80% of the insertions (78.5% for NA12878 and 78.3% for NA24385; **Figure 2** and **Figure S7**).

**Figure 2.**
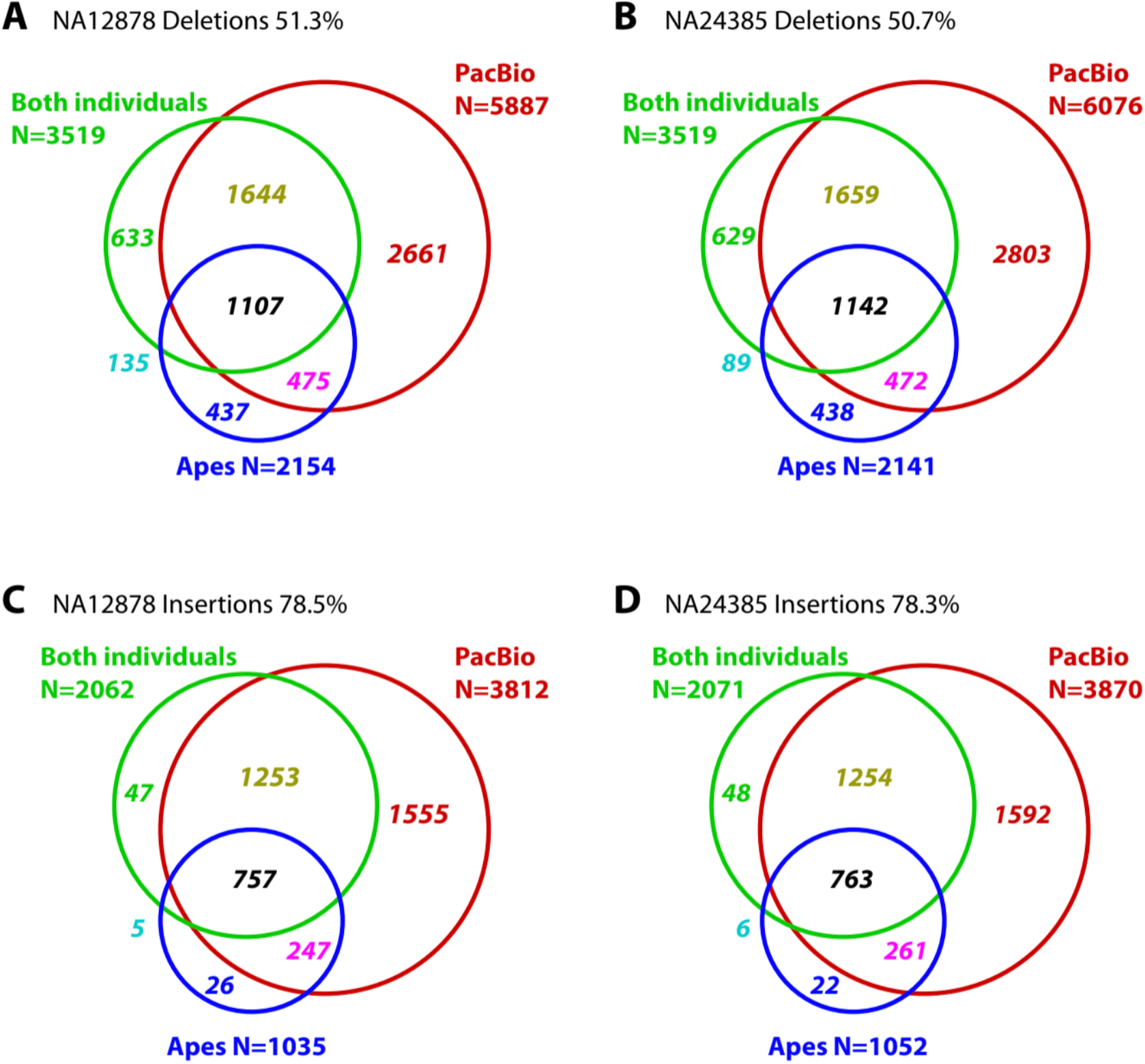
Three SV evaluation approaches as 1. overlap between NA12878 and NA24385 (Both individuals, green); 2. supported by any ape genome (Ape, blue); 3. supported by PacBio reads (PacBio, red). Numbers are SV counts.

Overlaps of calls between the two individuals or between one individual and an ape are likely to be highly specific, but not sensitive: specific because it is extraordinarily unlikely to produce the same SV twice in two independent hominid lineages; not sensitive because the two individuals do not share all variants, but rather a fraction that depends on population genetic parameters and stochasticity. The PacBio reads, by contrast, are derived from the same individual and are therefore expected to be both sensitive and specific. Indeed, PacBio reads validated the largest fraction of our SV calls compared to the other methods (**Figure 2** and **S7**). However, about 20% of deletions with support from apes, and about 18% of deletions with support from the other individual, were not validated by PacBio reads, suggesting that validation by PacBio is not fully sensitive either, and that some of the unvalidated deletion calls are in fact true positives. For insertions, the fraction of calls validated by the other individual but not by PacBio is considerably lower (ca 4%), consistent with the idea that insertion calls are more specific than deletion calls, as also suggested by their lower number.

We next investigated whether the type of sequence influenced the validation rate. Classification of insertions and deletions into Alu, non-Alu repetitive, and non-repetitive revealed considerably higher validation rates (by any of the aforementioned criteria) of Alu insertions than for the other two classes (**Figure 3** and **Figure S8**-**S9**). This is presumably because the assembly process is unlikely to produce a full-length Alu sequence erroneously, and so any insertion whose sequence matches an Alu is highly likely to be correct. Conversely, the fact that different assemblies produce a large number of unique Alu insertion calls that are likely correct again underscores that sensitivity of insertion detection is low, but specificity is high.

**Figure 3.**
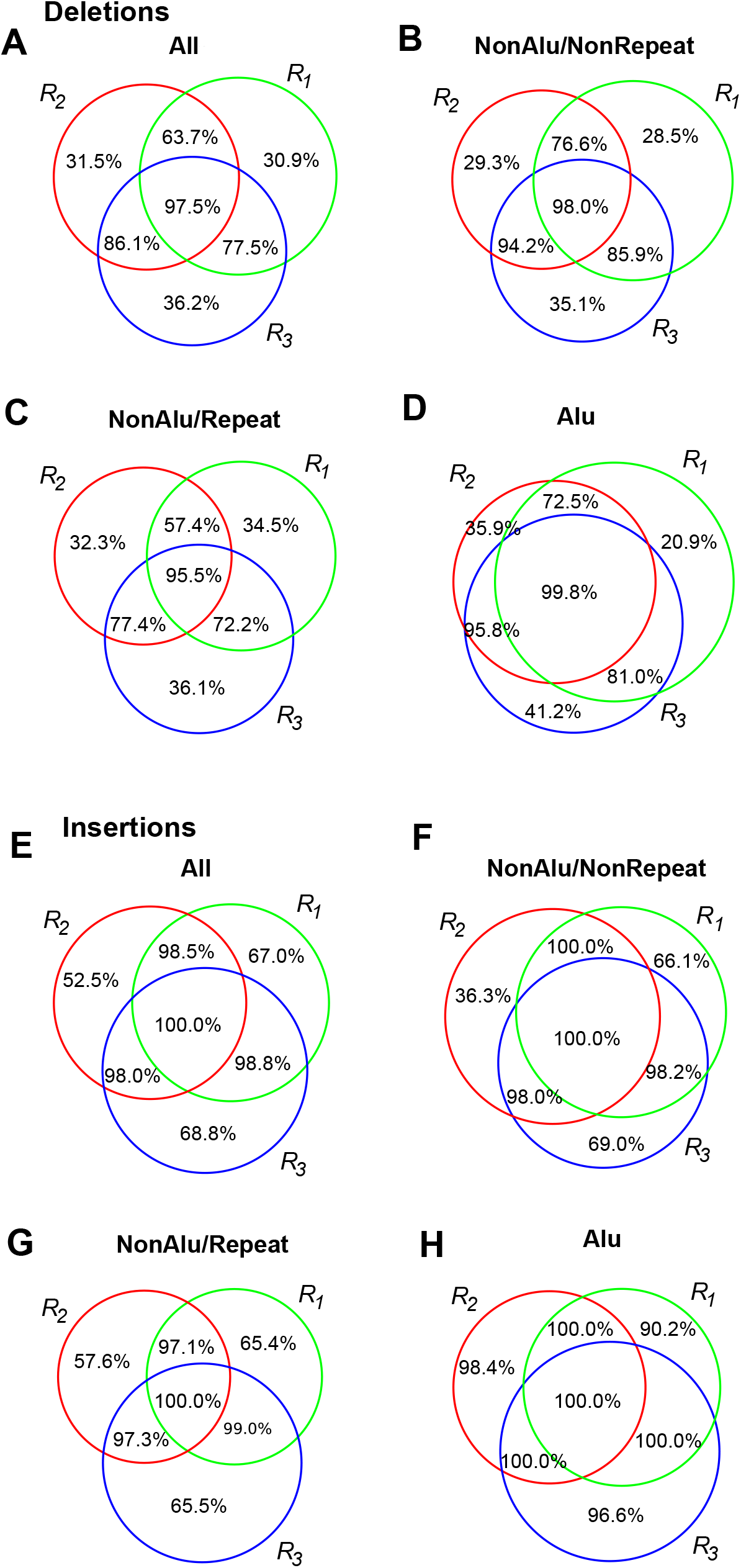
Sensitivities of deletions (A,B,C and D) and insertions (E, F, G and H) for the three libraries of NA12878. Percentages denote the proportion of SVs from assembly-based calls validated by any of the three evaluation approaches.

Finally, we examined whether the validation rate differed between unique SV calls or the shared ones among the assemblies from the same individual. As expected, the overall validation rate of SVs shared by all three libraries was greater than 95%, whereas unique SVs reached ca. 30% for deletions and ca. 50% for all insertions (**Figure 3**).

### Genotype accuracy and breakpoint precision of SV calls

To further evaluate assembly-based SV calls, we also assessed the accuracies of genotypes. As before, we validated unique and shared SV calls among the three libraries for each individual using PacBio reads. Overall, shared deletions reached above 68% genotype accuracy, with the subset that comprises Alus achieving 84%. Unique deletions reached above 40% accuracy. For insertions, accuracies for both shared and unique ones were significantly higher, above 92% and 75%, respectively. Shared Alu insertions achieved perfect accuracy (100%) (**Figure S10**-**S13**).

Finally, to assess the base-pair level accuracy of the SV breakpoints, we binned the SVs shared by both individuals based on their size differences between the two calls and evaluated their validation rates by either PacBio reads or the alignments to ape genomes. If the SVs were validated in both of the individuals, more than 80% of the deletions and 70% of the insertions had size differences smaller than 2bp. The rates were lower for calls not validated (60% for deletions, 40% for insertions; **Figure S14**).

### SV calls versus actual molecular mechanism

SVs are called ‘insertions’ or ‘deletions’ by comparison to the reference sequence, but that call does not necessarily reflect the actual molecular mechanism that gave rise to the SV: if the reference sequence carries the derived allele and our sequenced individual carries the ancestral state, the call is the opposite of the molecular mechanism. For 12,537 SVs, 1kb of flanking sequence (500bp on either side) could be aligned to at least one of their ape orthologs (**Methods**). On the basis of these alignments, assuming that the ape sequence represents the ancestral state, we thus classified each such SV as either a true insertion or a true deletion (**Figure 4A**). As expected from population genetic principles, a large fraction (37%) of deletion calls were in fact derived insertions, and half of called insertions were in fact deletions.

**Figure 4.**
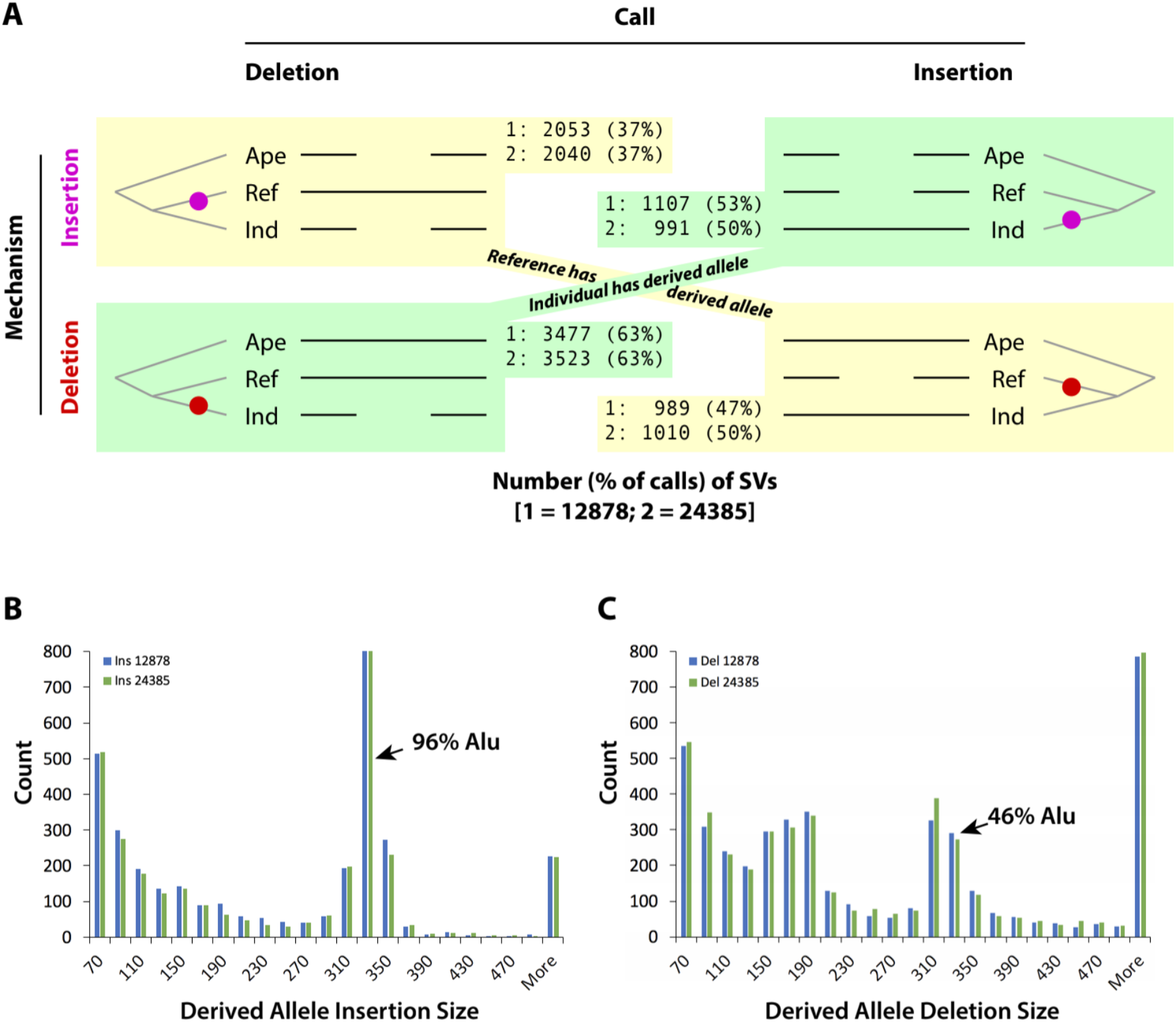
Classification of insertion and deletion calls into ancestral and derived state and inference of the originating molecular mechanism by comparison against ape genomes. A. Inference of derived allele and molecular mechanism by alignment to ape sequences; colored circle on tree denotes the lineage in which the mutation occurred; B. Derived allele insertion size distribution; C. Derived allele deletion size distribution.

Evidence that the derived allele actually reflects the molecular mechanism that initially generated the variant can be found in the size distribution of the events. Insertions (**Figure 4B**) follow an exponential dropoff in frequency as a function of size, with the major exception being a peak at 310-330 base pairs, in which 96% of insertions are full-length Alu sequence. By contrast, the deletion size distribution (**Figure 4C**) exhibits two regions of deviation from an exponential distribution, from ca. 110 to 150bp and from 290 to 330bp; the latter is somewhat enriched for Alu sequence, reflecting either (1) that we do not classify all called insertions correctly or (2) that there is some propensity for Alu elements to be deleted across their full lengths or nearly their full lengths. We also note that the vast majority of detected polymorphic L1 insertions were called as deletions in the assembled individual (i.e. the reference sequence carries the derived insertion allele), suggesting that SusperNova2 has a hard time assembling through young L1s that have not yet accumulated SNVs or other small variants.

### Ancient SVs

For 5,167 SVs, the two human sequences (reference and alternate allele plus 1kb flanking sequence as above) could be aligned to both orang and chimp orthologs. The vast majority of alignments were consistent between the two apes, supporting either the reference allele or the alternate allele as being ancestral. However, there were 225 events for which the chimp aligned to one allele, and the orang to the other (**Figure 5A**). Such inconsistencies can only be explained by two possibilities: (1) two independent insertions or deletions, one having occurred in one of the ape lineages, and another of the same sequence and coordinates generating the human derived allele; or (2), an ancient polymorphism that arose before our last common ancestor with chimp and that has been maintained in the human population since.

**Figure 5.**
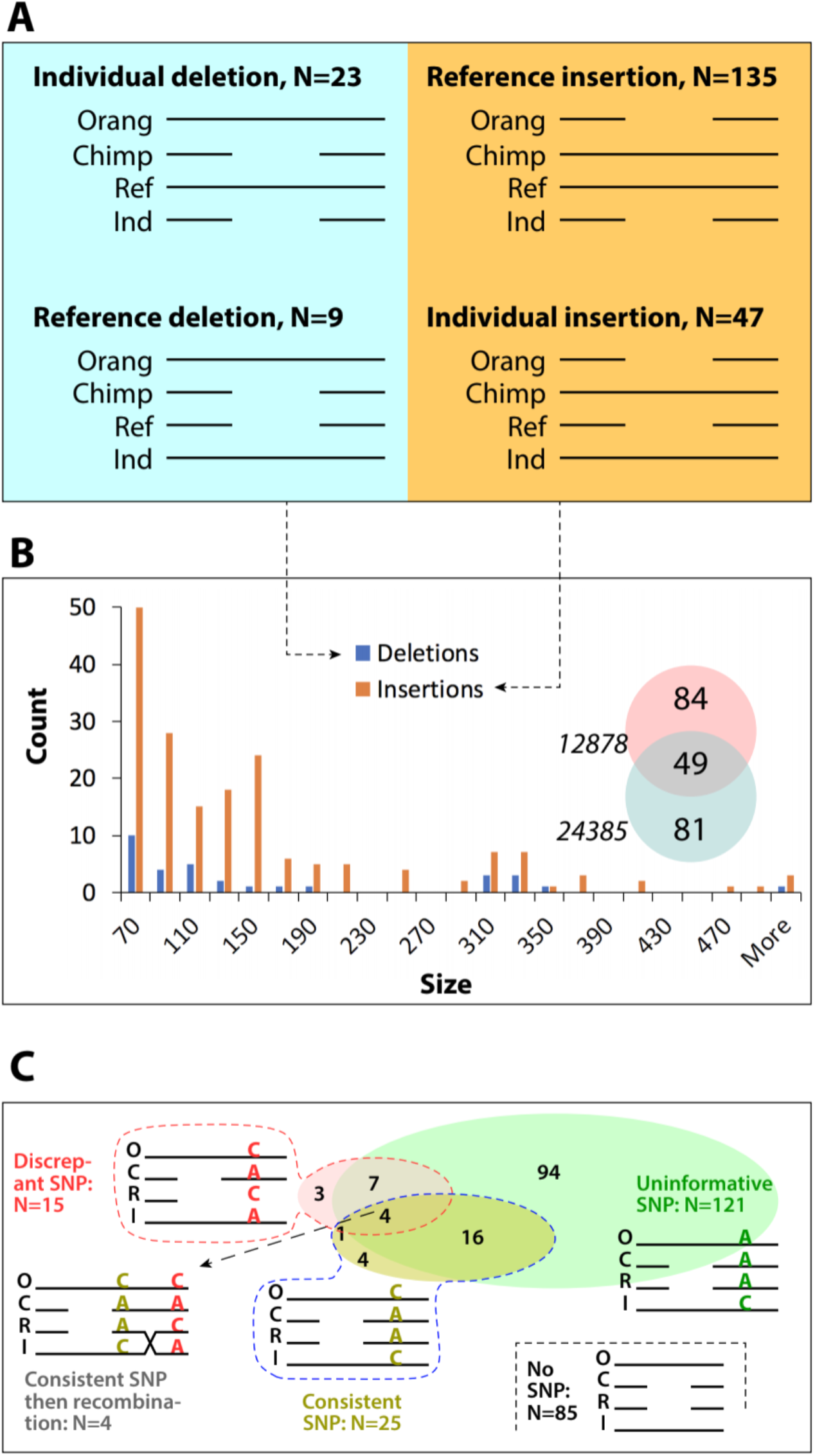
Ancient origin of SVs. A. The four cases in which orang matches one human allele and chimp the other, and their count in our dataset; B. Size distributions of the inferred 32 deletions and 182 insertions; Venn diagram indicates how many are shared between the two individuals and how many are unique to one of them; C. Phasing the SVs with closely linked SNVs; counts in Venn diagram indicate the number of each configuration.

To distinguish between these two possibilities, we proceeded as follows. SVs in our data sets that aligned to both chimp and orang occur approximately once per half-megabase (5,167 / length of genome covered in diploid contigs), and they are not clustered anywhere in the genome. The evolutionary distance between the apes and human is quite close, and while no models exist from which the probability of a hypothetical co-occurrence of SVs could be predicted, the proportion of such events in our data set (225 / 5,167 = 4%) seems quite high. To distinguish between the two possibilities, we constructed multiple sequence alignments among the four sequences and visually inspected each of them. For 214 events, we verified that the ape and human breakpoints precisely aligned and that the sequence of the ape SV was identical (excepting an occasional SNV or small indel) to that of the human allele. Size, sequence, and breakpoint locations of overlapping parallel events, by contrast, would be expected to be vary independently in humans and apes. We did not observe any such variance for the vast majority of the shared events, strongly suggesting that each SV has a single evolutionary origin and represents an ancient polymorphism maintained in our population since our last common ancestor with chimp.

Assuming that the orang sequence represents the ancestral state, we classified the SVs according to molecular mechanism, yielding 182 derived insertions and 32 derived deletions (**Figure 5A** and **5B**). This represents a highly significant (Chi-square test, P<10E-24) deviation from expectation (108 deletions, 106 insertions, based on their proportion in the set of 5,167 SVs that could be aligned to both apes). This deviation is consistent with the idea that insertion sequence is more likely than a deletion to produce evolutionary novelty and may be selected for. This finding represents indirect evidence for the selection (positive or balancing) that would be necessary to maintain these polymorphisms for such a long time.

Finally, the multiple alignments provided further opportunity to test the ancient polymorphism hypothesis by analysis of linked SNVs (**Figure 5C**). 129 alignments had at least one SNP in the 1kb of sequence surrounding the SVs; 94 of them were not informative, that is, both ape sequences had the same base, shared with either the reference or the individual. 25 alignments had at least one SNV that was in phase with the SV; 13 alignments had 5 or more phased SNVs. Curiously, 15 alignments had SNVs that were out of phase with the SV, and 5 of these also had at least one SNV that was in phase. Four of these 5 were arranged such that the SNVs with consistent phase were closer to the SV and the SNV with inconsistent phase was further away, suggesting that these four alignments capture not only ancient polymorphisms (SVs and SNVs) but also ancient recombination events between the consistent and the inconsistent SNVs. The considerable fraction of alignments that contain phased SNVs in the immediate vicinity of an SV is perhaps the strongest evidence in favor of the ancient polymorphism hypothesis.

## Discussion

Structural variants are abundant and important but require long-range information for their detection and so are not easily identified by standard (short-fragment) sequencing. We here explored the utility of assembly-based approaches for SV detection, specifically by using *de novo* assembly on the basis of 10x Genomics data. Our study demonstrates the promising future of assembly-based approaches to detect SVs in personal genomes, with reasonable sensitivity and genotype accuracy. Importantly, our pairwise-alignment based SV calls had remarkable breakpoint consistency and accuracy as evaluated by comparisons between the two individuals and with ape sequences.

### Diploid assembly and variant detection

In the context of diploid assembly, which is the natural approach for assembly of genomes of diploid organisms that harbor variation, the fraction of the genome that is assembled in a diploid state is a metric that needs to be carefully considered: it directly impacts variant discovery and genotyping, in that erroneously haploid regions will be missing all of their heterozygosity. The short input fragment length (*Wμ*_*FL*_ or *μ*_*FL*_) of *R*_1_ resulted in roughly 20% less of the genome in a diploid state (**Table 1** and **S1**, <60% vs <80%) compared to the other libraries of the same individual. As a consequence, there were fewer SNV and SV calls in the analyses involving *R*_1_ (**Table S2**).

Sensitivity of SNV detection is naturally limited by the fraction of the genome that is covered by the assembly, and genotype accuracy evaluation is limited to the fraction of the assembly that is in a diploid state. Overall sensitivity of assembly-based calls is ca. 90% of that of mapping-based SNV calls and incorrect call rates in high-confidence regions of GIAB are also higher than that in mapping-based calls. We conclude that at this point, assembly-based SNV calls from SuperNova2 were not competitive with barcode-aware read-mapping approaches. However, we note that this is not a compromise as exactly the same sequence data can be used for SNV detection (via barcode-aware mapping) and SV detection (via assembly). We estimate that the cost increase over standard Illumina sequencing is about 2x, given the 10X prep cost and the higher level of sequence coverage required. There may be many applications for which this combination of excellent SNV detection (via barcode-aware read-mapping) and highly precise SV discovery (via assembly), achieved by the same data set, is worth the cost.

### Importance for *de novo* assembly-based SV detection

Our study highlights two concepts that are important for SV science. The first is that the variation call that is based on comparison to reference is not the same as the allelic origin of the variant. Molecularly, that allelic origin is also the mechanism that gave rise to the variant as the initial single mutation that arose in an ancestral individual’s germline. In our individuals, very large fractions of deletion calls were actually insertions, and vice versa, as expected and as illustrated with hundreds of Alu insertions. The second concept is that there may be many more regions than previously thought in which heterozygosity has been maintained in our lineage since before our last common ancestor with chimp. Our results in this regard support the idea that there is distinct value in assembly-based approaches for determining SVs in large numbers of individuals for population genetic questions as well.

## Methods

### DNA extraction, library construction and sequencing

For library *L*_1_, genomic DNA was extracted from ca. 1 million cultured 12878 cells using the Gentra Puregene Blood Kit following manufacturer’s instructions (Qiagen, Cat. No 158467). To generate longer DNA fragments (W*μ*_*FL*_=150kb and longer) for *L*_2_ to *L*_6_, a modified protocol for DNA extraction was applied. Two-hundred thousand NA12878 or NA24385 cells of fresh culture were added to 1mL cold 1x PBS in a 1.5 ml tube and centrifuged for 5 minutes at 300g. The cell pellets were completely resuspended in the residual supernatant by vortexing and then lysed by adding 200ul Cell Lysis Solution and 1ul of RNaseA Solution (Qiagen, Cat. No 158467), mixing by gentle inversion, and incubating at 37°C for 15-30 minutes. This cell lysis solution is used immediately as input for the 10x Chromium prep (ChromiumTM Genome Library & Gel Bead Kit v2, PN-120258; ChromiumTM i7 Multiplex Kit, PN-120262). Fragment size of the input DNA can be controlled by gentle handling during lysis and DNA preparation for Chromium. Different amounts of input DNA (between 1.25 and 4 ng) were used to generate libraries with different *C*_*F*_. The Chromium Controller was operated and the GEM prep was performed as instructed by the manufacturer. Individual libraries were then constructed by end repairing, A-tailing, adapter ligation and PCR amplification. Each library was sequenced with three lanes of paired-end 150bp runs on the Illumina HiSeqX instrument to obtain high genomic coverage.

### *De novo* diploid assembly

Scaffolds were generated by the “pseudohap2” output style of SuperNova2, which explicitly generated scaffolds for two haplotypes, simultaneously. Pairs of scaffolds were extracted as the two haplotypes from the SuperNova2 megabubble structures if they shared the same start and end nodes in the assembly graph. Diploid contigs were generated by breaking the candidate scaffolds at the sequences with least 10 consecutive ‘N’s and were aligned to human reference genome (hg 38) by Minimap2^46^. The genome was split into 500bp windows and diploid regions were defined as the maximum extent of successive windows covered by two contigs, each from one haplotype.

### SNV and structural variant calls from diploid contigs

We used Paftools (https://github.com/lh3/minimap2/tree/master/misc) to identify SNVs and SVs no shorter than 50bp from the CS tags generated by Minimap2 alignment. A valid variant was covered by exactly two contigs with mapping quality larger than 20, each from one haplotype. SVs were called as homozygous if the calls from the two allelic contigs were overlapping. SVs were considered shared among assemblies from the same individual if there was any overlap in coordinates.

### Validation of SNV calls

We validated SNVs by comparison with the ‘gold standard’ GIAB SNV call set (NA12878: ftp://ftp-trace.ncbi.nlm.nih.gov/giab/ftp/release/NA12878_HG001/latest/GRCh38/, NA24385: ftp://ftp-trace.ncbi.nlm.nih.gov/giab/ftp/release/AshkenazimTrio/HG002_NA24385_son/latest/GRCh38/). Any SNV calls were removed if they are outside of GIAB high-confidence regions or diploid regions. The SNVs from barcode-aware read-mapping were generated by freebayes (https://github.com/ekg/freebayes) from the alignments of Lariat ^47^.

### Validation of SV calls

SVs were examined by three approaches: 1. We applied svviz2 ^45^ to analyze PacBio reads from NA12878 (ftp://ftp-trace.ncbi.nlm.nih.gov/giab/ftp/data/NA12878/NA12878_PacBio_MtSinai/) and NA24385 (ftp://ftp-trace.ncbi.nlm.nih.gov/giab/ftp/data/AshkenazimTrio/HG002_NA24385_son/PacBio_MtSinai_NIST/). svviz2 aligned and compared the PacBio reads to the reference sequence and the reconstructed alternative allele of candidate SVs. Genotypes 0/1 and 1/1 confirmed our SV calls; genotypes were also used to evaluate the genotype accuracy in the validated call set; 2. We identified SVs called in both NA12878 and NA24385 and considered them reciprocally validated if their coordinates differed by fewer than 20bp. We only considered the existence of SVs regardless of their genotype concordance. The complete set of SVs for each sample was the union of calls of the three libraries; 3. We aligned each SV and 500 bp flanking sequence on either side from the involved contigs to their chimp (reference genome Pan_tro_3.0) and orang (reference genome PPYG2) orthologs. We defined the distance between the end of the left flanking sequence and the start of the right flanking sequence as *dis*(align). For deletions, if *dis*(align) was smaller than 2bp, then the derived allele was recognized as an insertion carried by the reference genome; if *dis*(align) was between 0.9 to 1.1 times of the SV length, then the derived allele was recognized as a deletion in the individual’s genome. For insertions, if *dis*(align) was smaller than 2bp, then the derived allele was recognized as an insertion carried by the target genome; if *dis*(align) was between 0.9 to 1.1 times of the SV length, then the derived allele was recognized as a deletion carried by the reference genome.

### Annotation of SV sequence

Deletions and insertions were annotated as Alu sequences if they were between 250bp to 350bp long and could be uniquely aligned to the Alu consensus sequence. We used Tandem Repeats Finder ^48^ to annotate tandem repeats.

### Multiple sequence alignment to detect ancient polymorphism

We produced the four-way multiple sequence alignments using Muscle ^49^ from the SVs where orang and chimp differed in matching the reference sequence or the alternate allele. The sequences were 1. human reference sequence; 2 assembled target sequence; 3. orangutan (version) reference sequence; and 4. chimpanzee reference sequence (version). We then examined all such alignments to verify that the SV sequence was orthologous and that the breakpoints were identical.

## Data access

Assemblies are currently available at http://mendel.stanford.edu/supplementarydata/zhang_SN2_2019. We will be submitting raw sequence data and assemblies to NCBI’s SRA and Assembly databases.

## Acknowledgements

This research was supported by training and research grants from the National Institute of Standards and Technology. We would like to thank Justin Zook, Marc Salit, Alex Bishara, Noah Spies, Nancy Hansen, David Jaffe, and Deanna Church for informative discussions.

