## Supplementary files for "*De novo* diploid genome assembly for genome-wide structural variant detection"

### Supplementary Tables

| Library | Sample id | Raw coverage (X) | $\mu_{FL}$/W$\mu_{FL}$ (kb) | PCR duplication (%) | $C_F$ (X) | $C_R$ (X) |
|---|---|---|---|---|---|---|
| $L_1$ | NA12878 | 192 | 24.0/41.1 | 10.92 | 334 | 0.27 |
| $L_2$ | NA12878 | 103 | 79.0/304.3 | 19.97 | 123.2 | 0.41 |
| $L_3$ | NA12878 | 106 | 99.2/214.5 | 11.09 | 958.7 | 0.07 |
| $L_4$ | NA24385 | 117 | 92.1/216.9 | 10.88 | 1504.6 | 0.05 |
| $L_5$ | NA24385 | 100 | 120.8/267.4 | 18.51 | 208.4 | 0.25 |
| $L_6$ | NA24385 | 100 | 64.2/151.7 | 12.39 | 803.3 | 0.08 |

**Table S1.** Parameters of 10x libraries prepared for NA12878 and NA24385.

| Library | SNVs | Deletions>50bp | Insertions>50bp |
|---|---|---|---|
| $R_1$ | 2,635,173 | 7,224 | 2,350 |
| $R_2$ | 2,820,432 | 6,604 | 2,650 |
| $R_3$ | 3,107,823 | 7,005 | 3,002 |
| $R_4$ | 2,900,187 | 6,918 | 2,779 |
| $R_5$ | 2,995,409 | 8,123 | 2,742 |
| $R_6$ | 3,100,365 | 7,359 | 3,117 |

**Table S2.** The number of SNVs and SVs in diploid regions detected by SuperNova2.

| Library | Sample | Total SNVs from assembly | Overlap with any other sets | Overlap of all the three sets |
|---|---|---|---|---|
| $R_1$ | NA12878 | 2,635,173 | 2,455,466 (93.2%) | 1,999,330 (75.9%) |
| $R_2$ | NA12878 | 2,820,432 | 2,606,749 (92.4%) | 1,999,330 (70.9%) |
| $R_3$ | NA12878 | 3,107,823 | 2,887,671 (92.9%) | 1,999,330 (64.3%) |
| $R_4$ | NA24385 | 2,900,187 | 2,800,750 (96.6%) | 2,436,887 (84.0%) |
| $R_5$ | NA24385 | 2,995,409 | 2,759,313 (92.1%) | 2,436,887 (81.4%) |
| $R_6$ | NA24385 | 3,100,365 | 2,931,116 (94.5%) | 2,436,887 (78.6%) |

**Table S3.** The number and proportion of shared SNVs in the three libraries for each individual.

| Library | Sample | Total SNVs shared by three sets | Set unique genotype | Not concordant by all the three sets |
|---|---|---|---|---|
| $R_1$ | NA12878 | 1,999,330 | 49,673 (2.5%) | 191 (0.0096%) |
| $R_2$ | NA12878 | 1,999,330 | 94,005(4.7%) | 191 (0.0096%) |
| $R_3$ | NA12878 | 1,999,330 | 23,244 (1.2%) | 191 (0.0096%) |
| $R_4$ | NA24385 | 2,436,887 | 25,173 (1.0%) | 162 (0.0066%) |
| $R_5$ | NA24385 | 2,436,887 | 110,188 (4.5%) | 162 (0.0066%) |
| $R_6$ | NA24385 | 2,436,887 | 23,179(1.0%) | 162 (0.0066%) |

**Table S4.** SNV genotype concordance of the three libraries for each individual.

| Library | Total SNV from assembly-based calls | Proportion of SNVs in mapping-based calls | Genotype concordance |
|---|---|---|---|
| $R_1$ | 2,635,173 | 92.4% | 99.8% |
| $R_2$ | 2,820,432 | 92.8% | 99.8% |
| $R_3$ | 3,107,823 | 93.6% | 99.8% |
| $R_4$ | 2,900,187 | 96.5% | 99.8% |
| $R_5$ | 2,995,409 | 95.1% | 99.8% |
| $R_6$ | 3,100,365 | 96.3% | 99.8% |

**Table S5.** SNVs sensitivity and genotype concordance between assembly-based and mapping-based SNV calls.

| Library | Overlap with GC | FP | FN | Genotype concordance | Het to Hom | Hom to Het |
|---|---|---|---|---|---|---|
| $R_1$ | 2,150,961 | 8,009 (3.59%) | 133,746 (5.85%) | 2,072,862 (96.37%) | 77,973 (3.63%) | 126 (0.0058%) |
| $R_2$ | 2,322,370 | 96,729 (4.00%) | 171,071 (6.86%) | 2,210,351 (95.18%) | 111,757 (4.81%) | 262 (0.011%) |
| $R_3$ | 2,568,775 | 93,618 (3.52%) | 102,443 (3.84%) | 2,574,889 (98.63%) | 35,069 (1.37%) | 214 (0.0083%) |
| $R_4$ | 2,252,797 | 20,699 (0.91%) | 49,784 (2.16%) | 2,218,744 (98.49%) | 33.767 (1.50%) | 286 (0.013%) |
| $R_5$ | 2,286,108 | 38,240 (1.65%) | 149,517 (6.14%) | 2,173,945 (95.09%) | 111,857 (4.89%) | 306 (0.013%) |
| $R_6$ | 2,393,193 | 22,349 (0.93%) | 53,403 (2.18%) | 2,357,389 (98.50%) | 35,473 (1.48%) | 331 (0.014%) |

**Table S6.** Sensitivity and genotype accuracy of SNVs from assembly-based calls by comparing with 'gold standard' GIAB call set (GC). FP: False Positives; FN: False Negatives; Het to Hom: heterozygous SNVs in GC that were miscalled as homozygosity; Hom to Het: homozygous SNVs in GC that were miscalled as heterozygosity.

#### Supplementary Figure

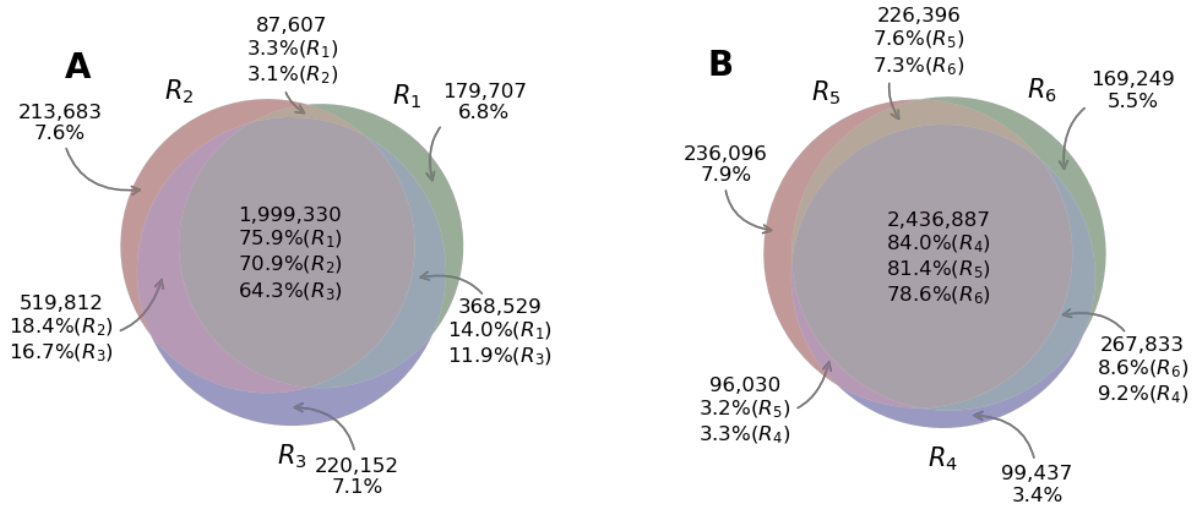


**Figure S1.** Overlaps of SNV calls in three libraries for each individual. (A) NA12878; (B) NA24385

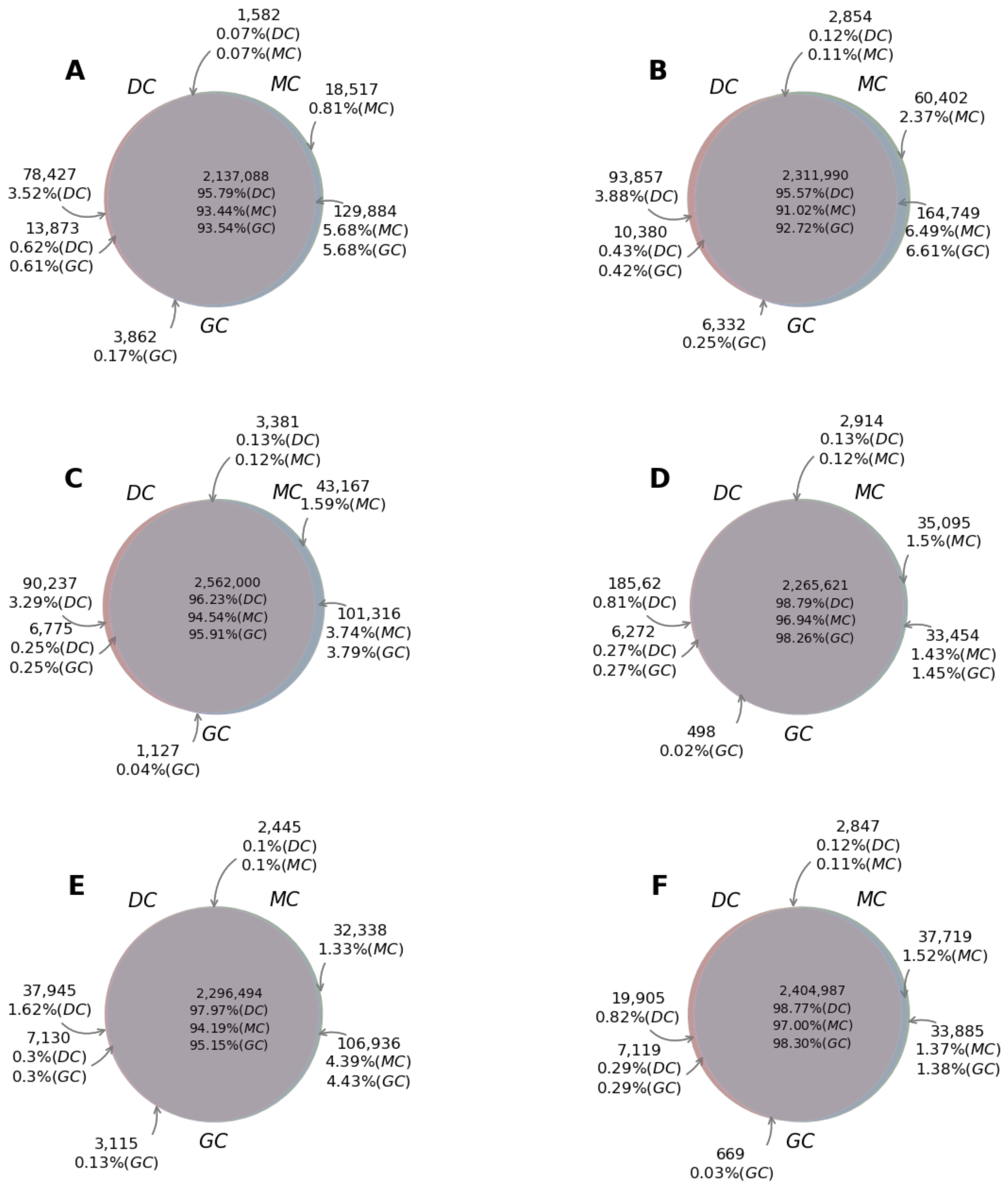


**Figure S2.** Overlap of SNV calls from DC (assembly-based calls), MC (mapping-based calls) and GC ('gold standard' GIAB call set). A: $R_1$ B: $R_2$, C: $R_3$, D: $R_4$, E: $R_5$, F: $R_6$. Counts and percentages are SNV counts and the fractions in each call set, respectively.

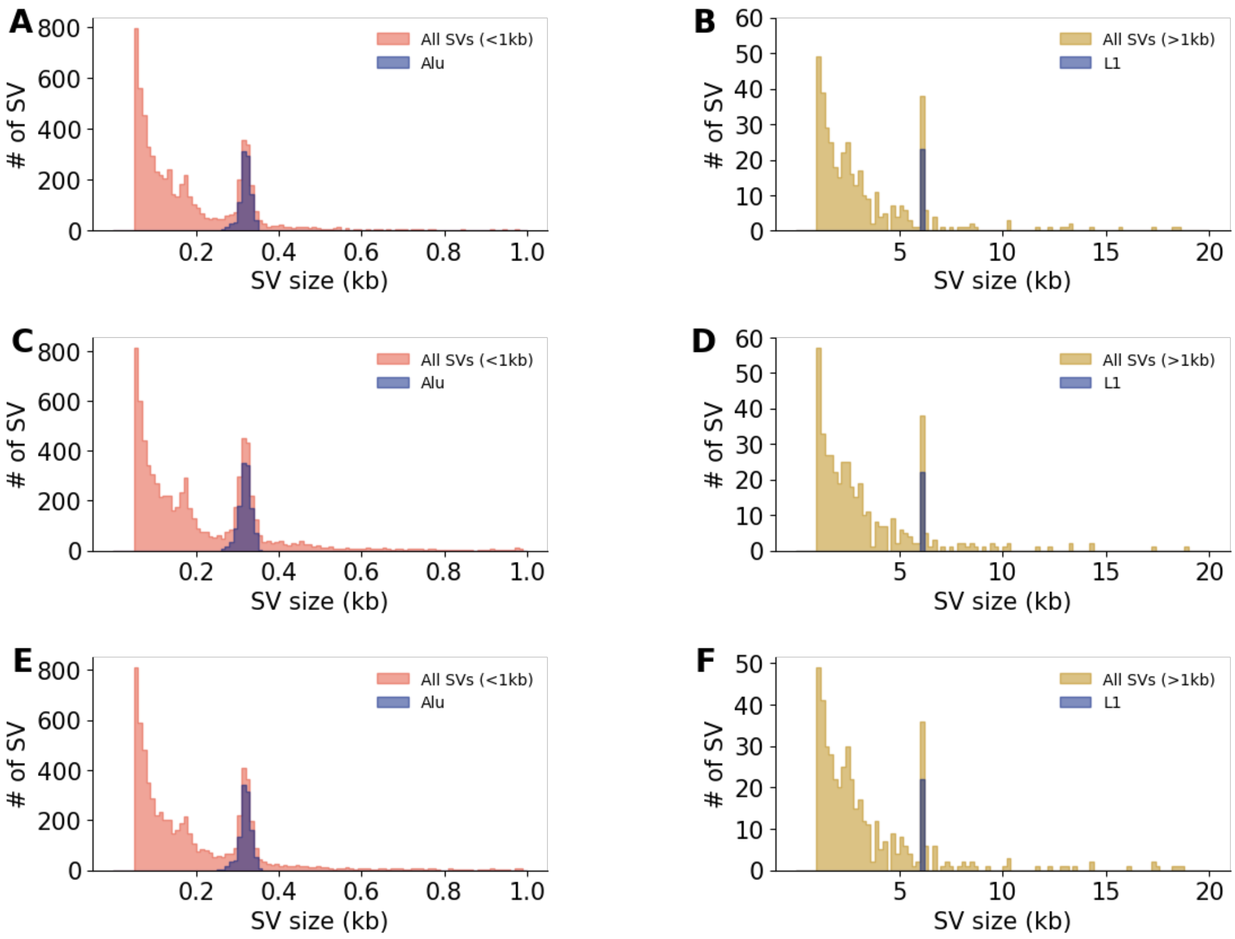


**Figure S3.** Deletion size distributions of NA24385 from $R_4$ (A and B), $R_5$ (C and D) and $R_6$ (**E** and **F**).

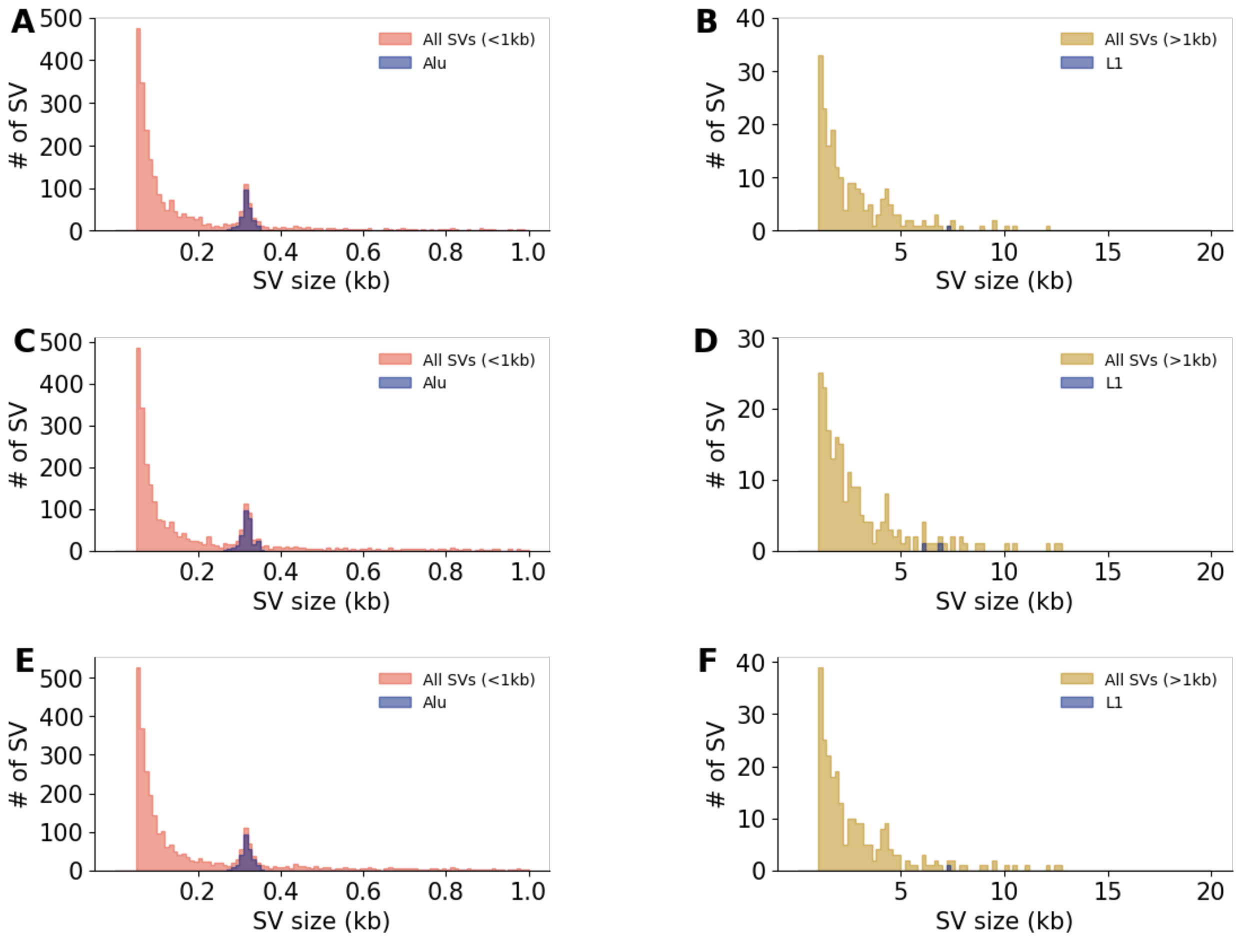


**Figure S4.** Insertion size distributions of NA24385 from $R_4$ (A and B), $R_5$ (C and D) and $R_6$ (E and F).

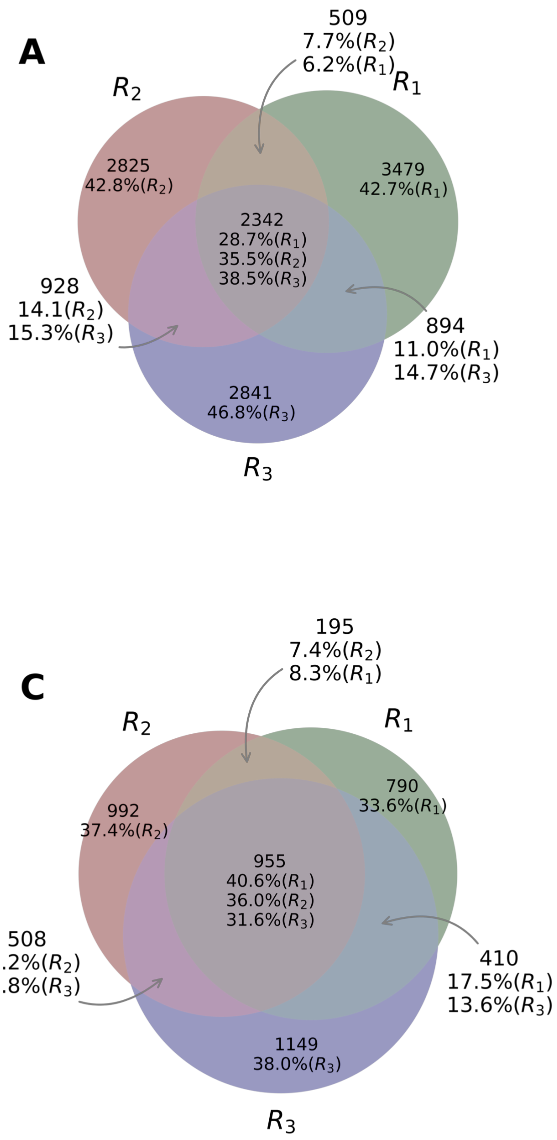


**Figure S5.** Overlap of deletions and insertions for three libraries from each individual detected by SuperNova2. A. Deletions from NA12878; B. Deletions from NA24385; C. Insertions from NA12878; D. Insertions from NA24385. Counts and percentages are SNV counts and the fractions in each call set, respectively.

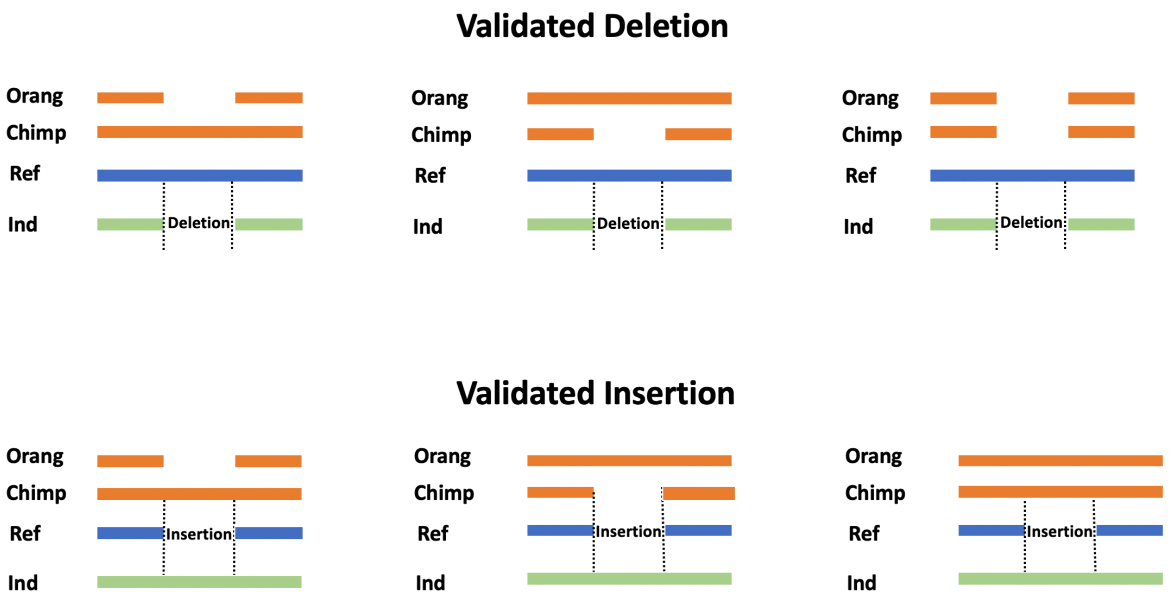


**Figure S6.** Evaluation of deletions and insertions by alignment to ape genomes.

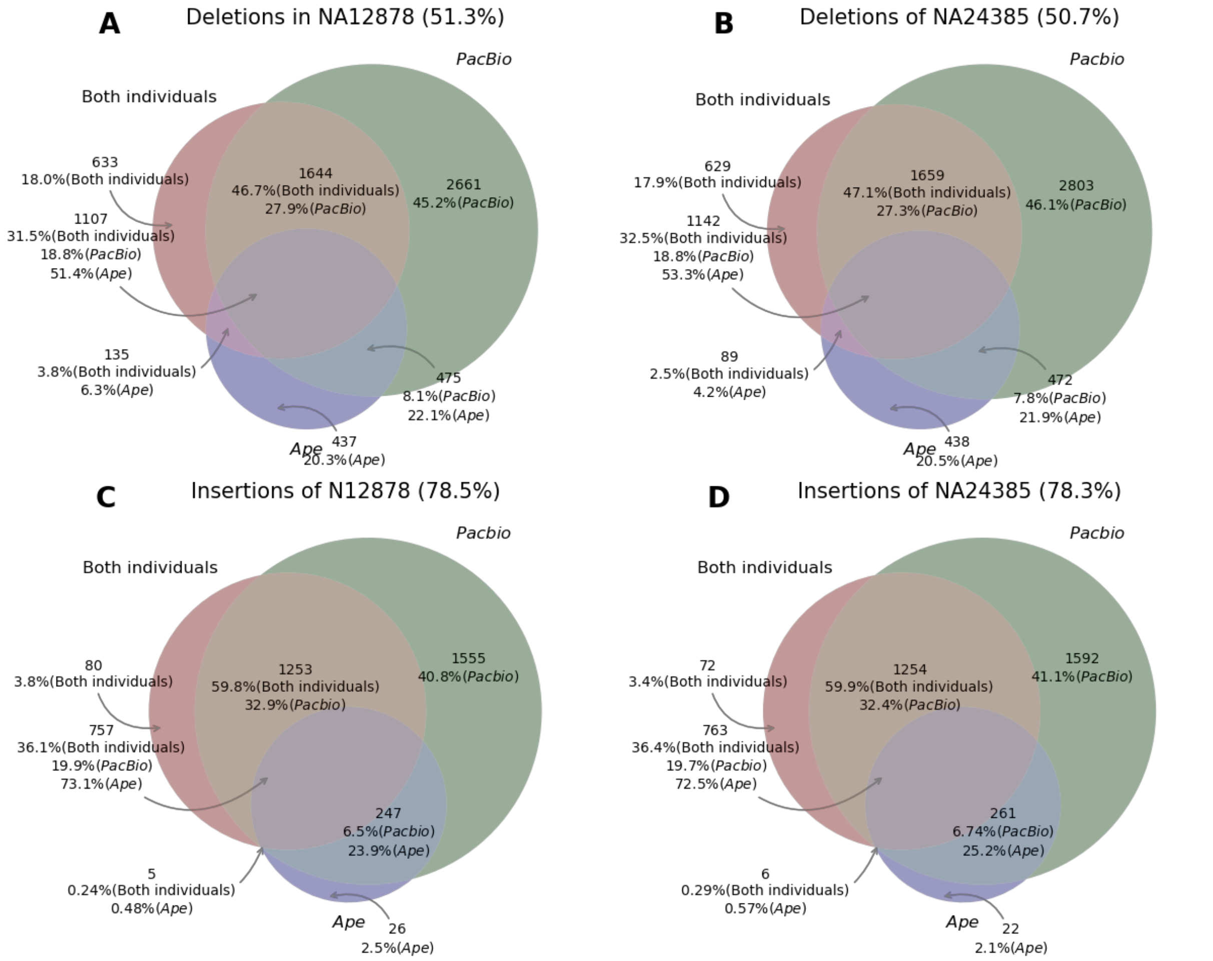


**Figure S7.** Comparison of three SV evaluation approaches (with percentage) as 1. overlap between NA12878 and NA24385 (Both individuals, red); 2. supported by any Ape genomes (Ape, blue); 3. supported by PacBio reads (PacBio, green). The numbers are SV counts shown in **Figure 5**. Percentages represent the fractions of validated calls in the corresponding validation approaches.

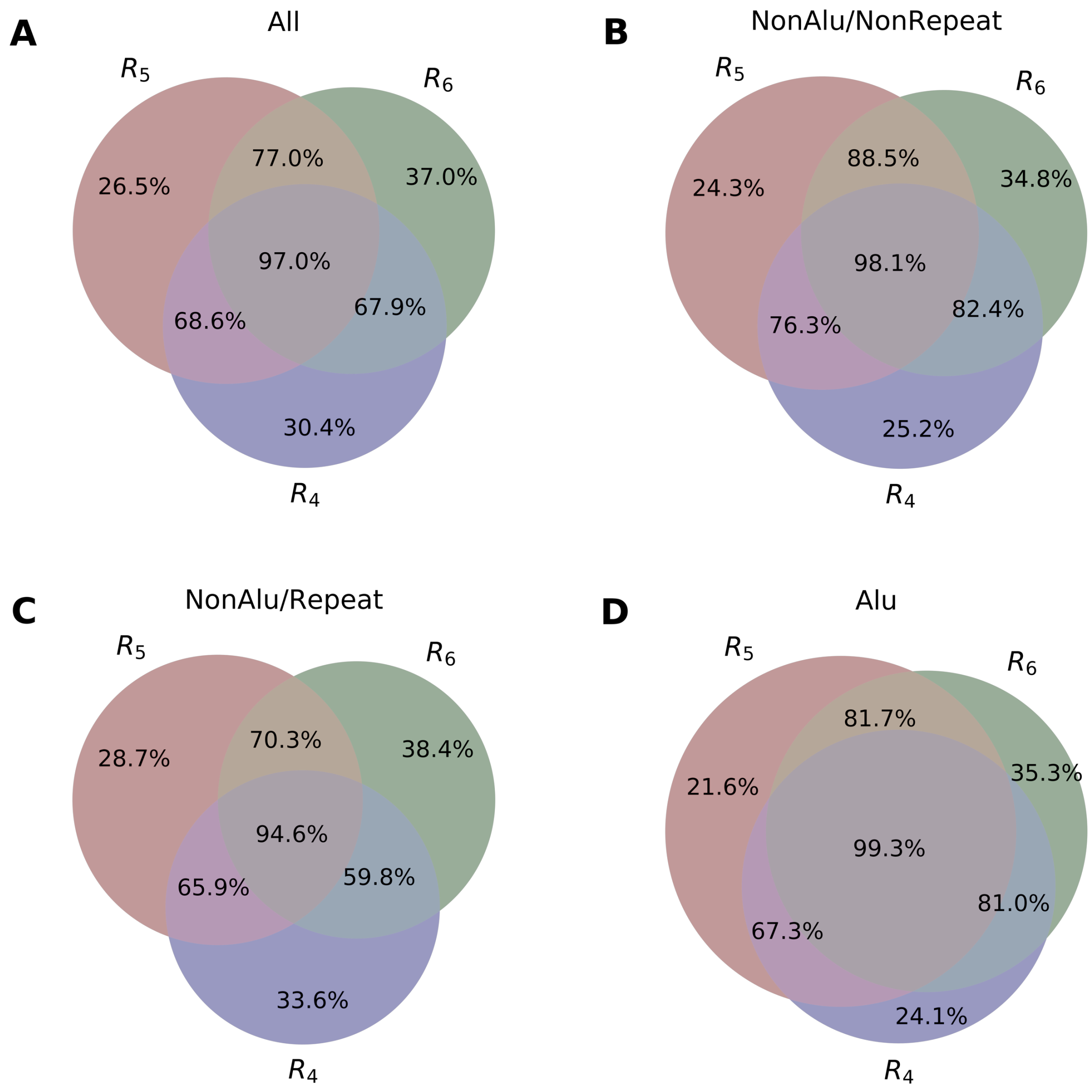


**Figure S8.** Deletion sensitivity of the three libraries for NA24385. The percentages denote the proportion of SVs from assembly-based calls that validated by any of the three approaches.

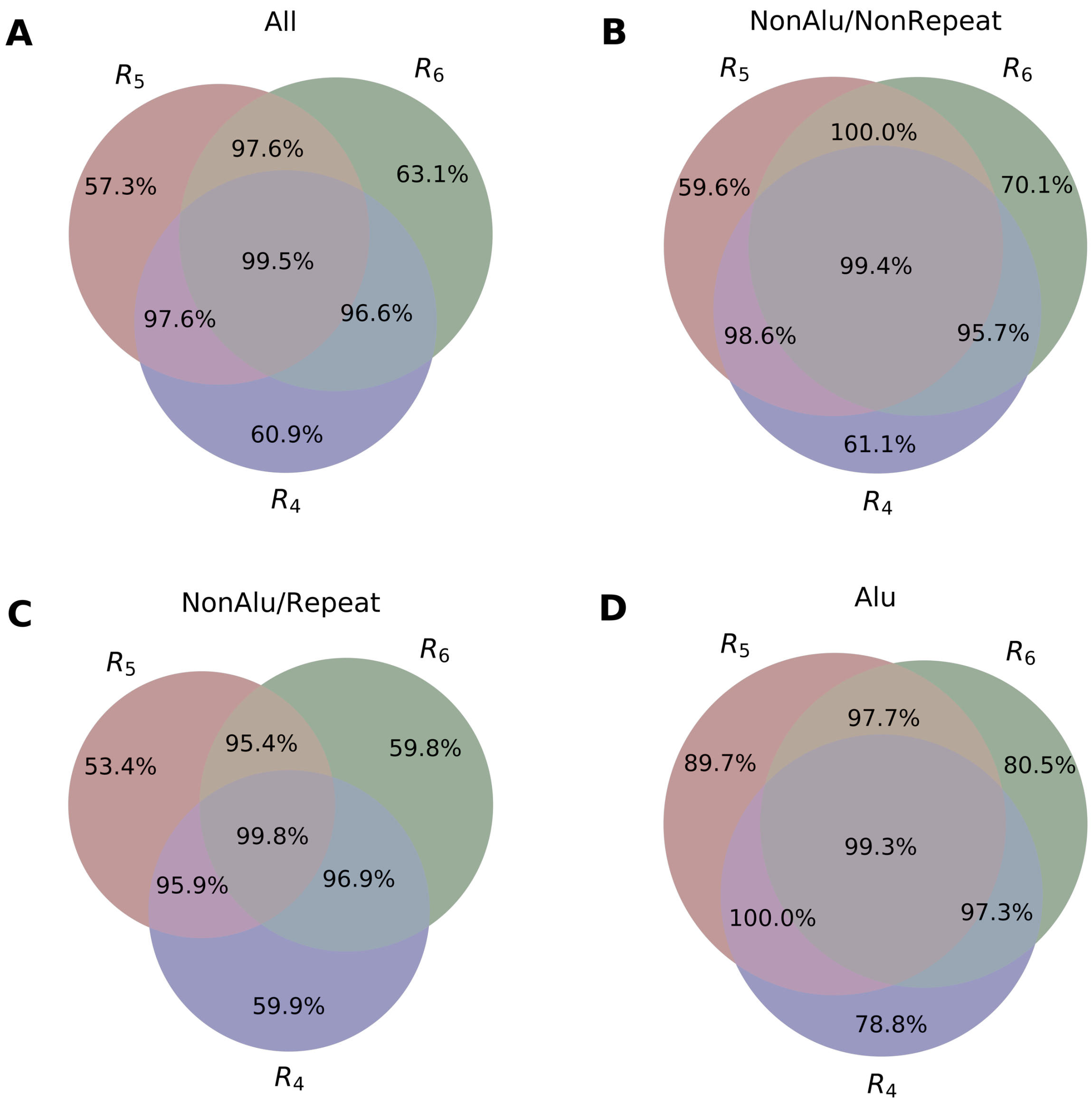


**Figure S9.** Insertion sensitivity of the three libraries for NA24385. The percentages denote the proportion of SVs from assembly-based calls that validated by any of the three approaches.

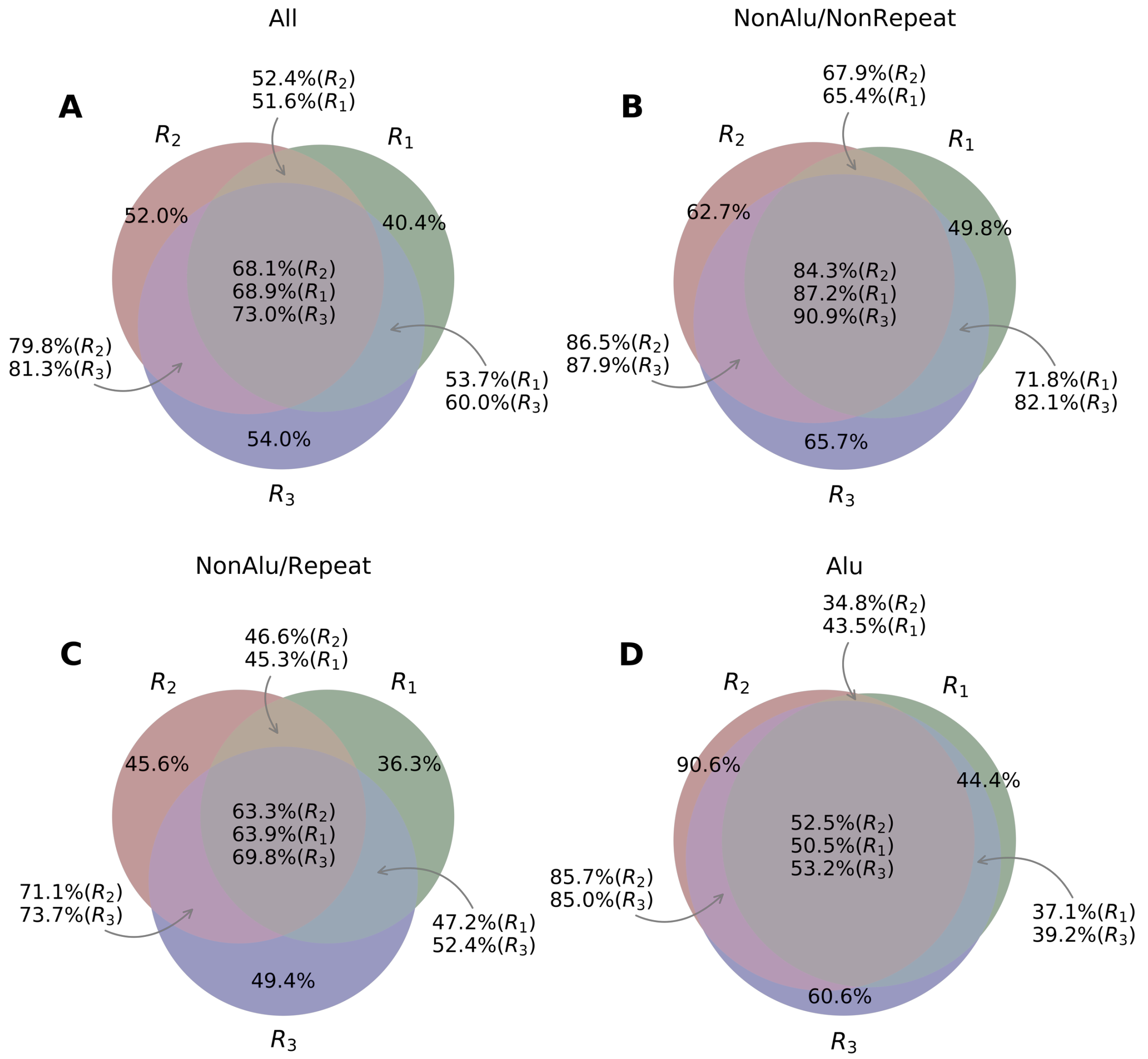


**Figure S10.** Genotype accuracy of deletions in NA12878 evaluated by PacBio reads. The percentages denote the proportion of SVs have consistent genotypes with PacBio reads.

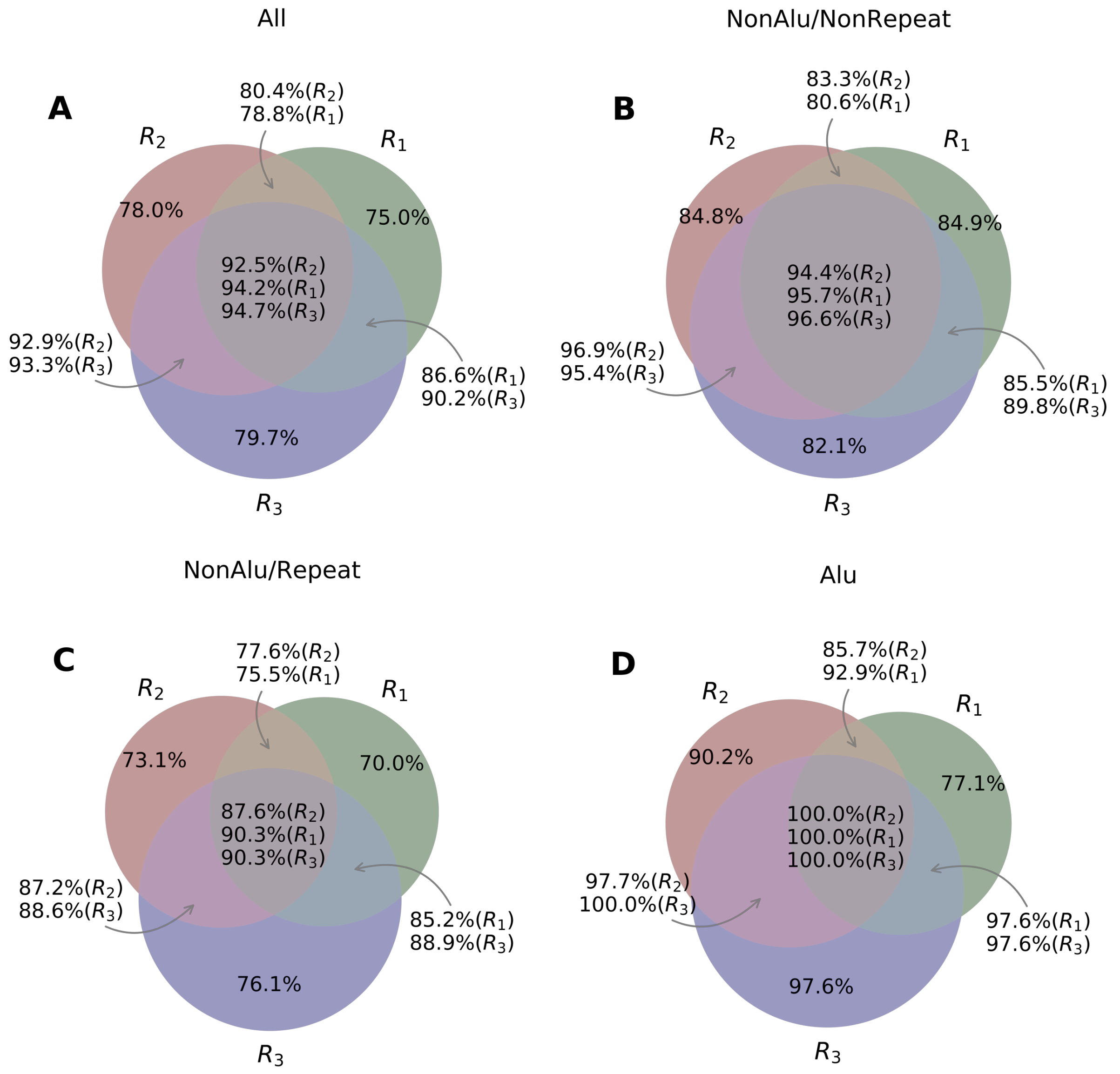


**Figure S11.** Genotype accuracy of insertions in NA12878 evaluated by PacBio reads. The percentages denote the proportion of SVs have consistent genotypes with PacBio reads.

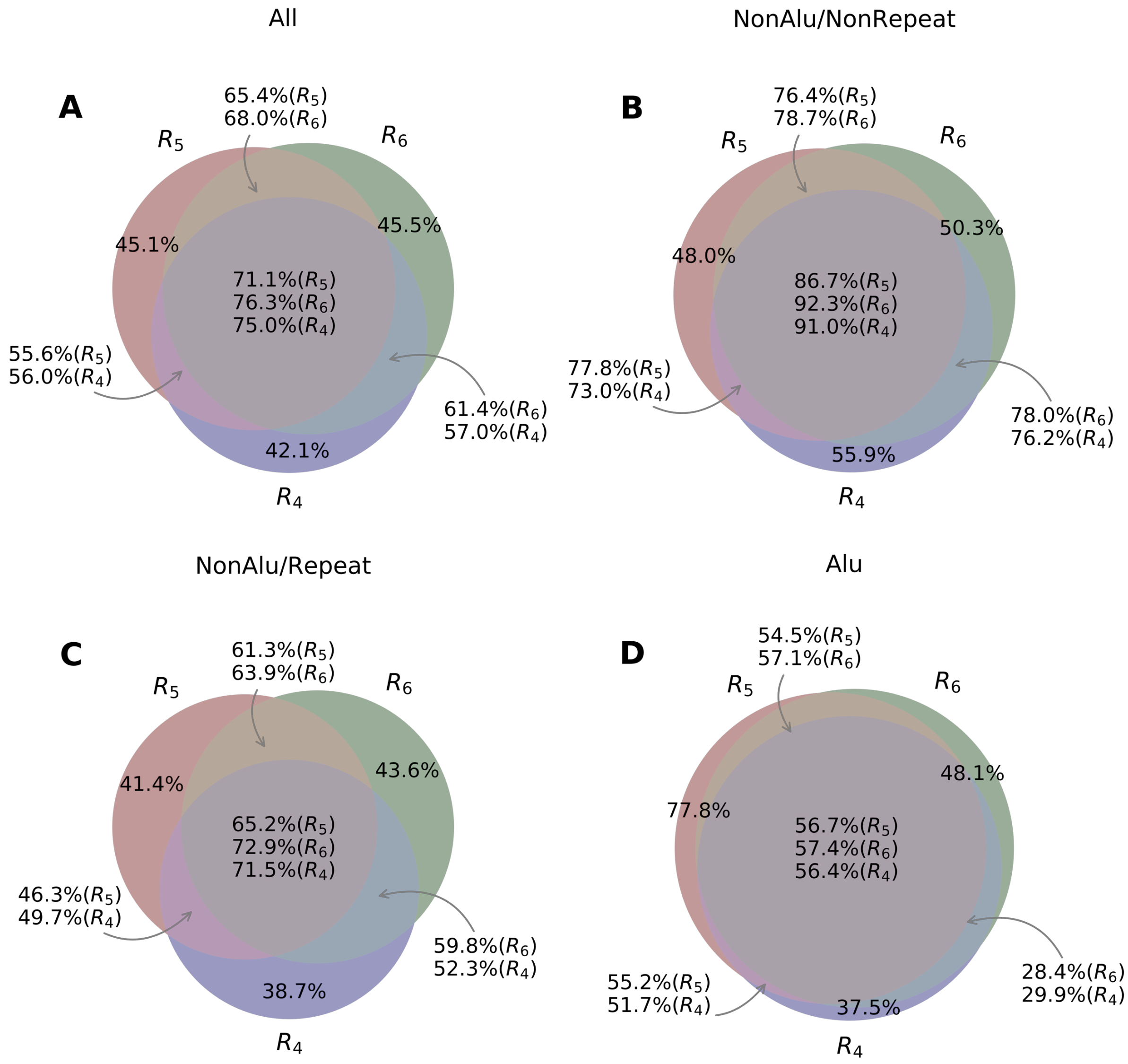


**Figure S12.** Genotype accuracy of deletions in NA24385 evaluated by PacBio reads. The percentages denote the proportion of SVs have consistent genotypes with PacBio reads.

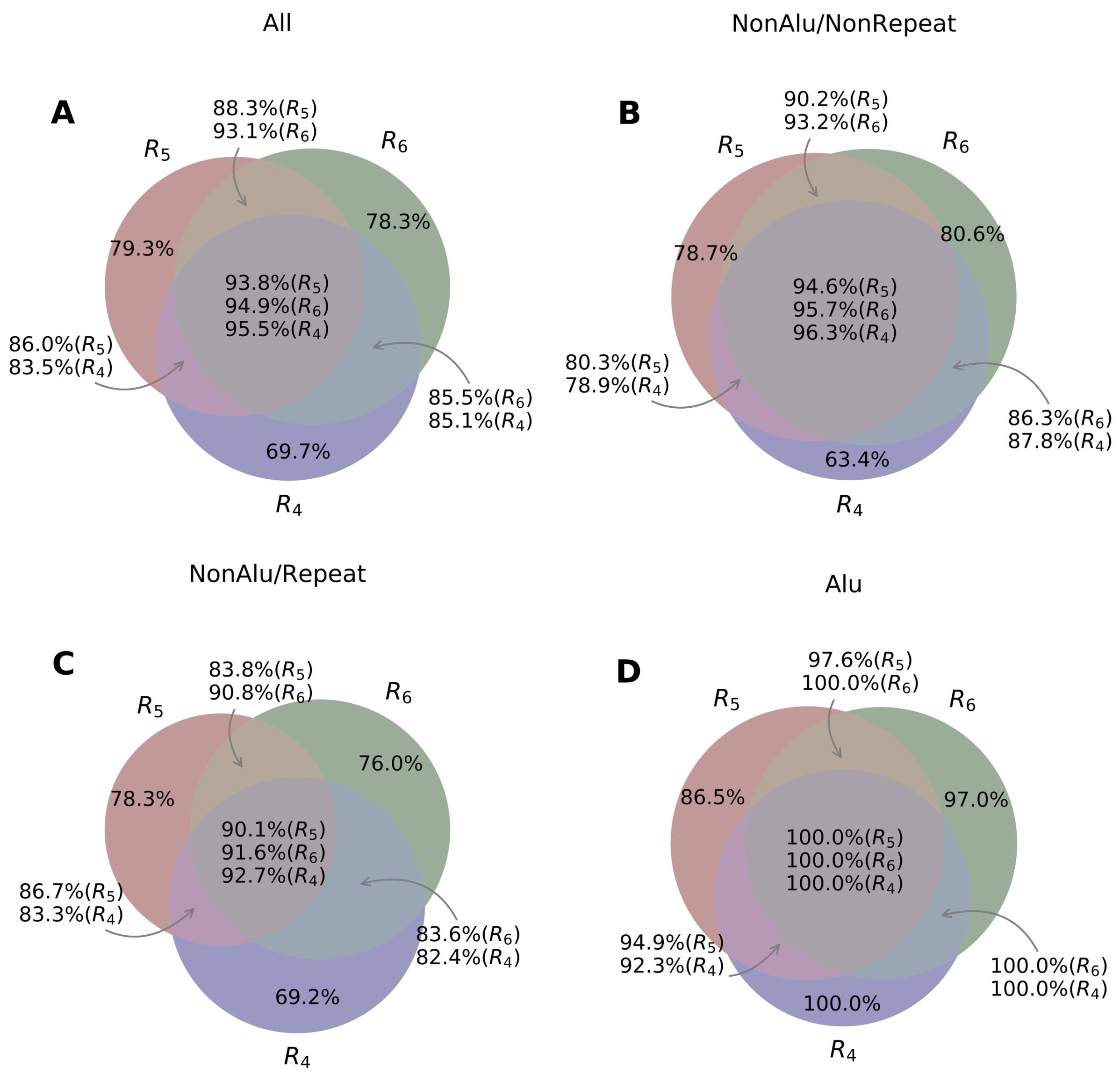


**Figure S13.** Genotype accuracy of insertions in NA24385 evaluated by PacBio reads. The percentages denote the proportion of SVs have consistent genotypes with PacBio reads.

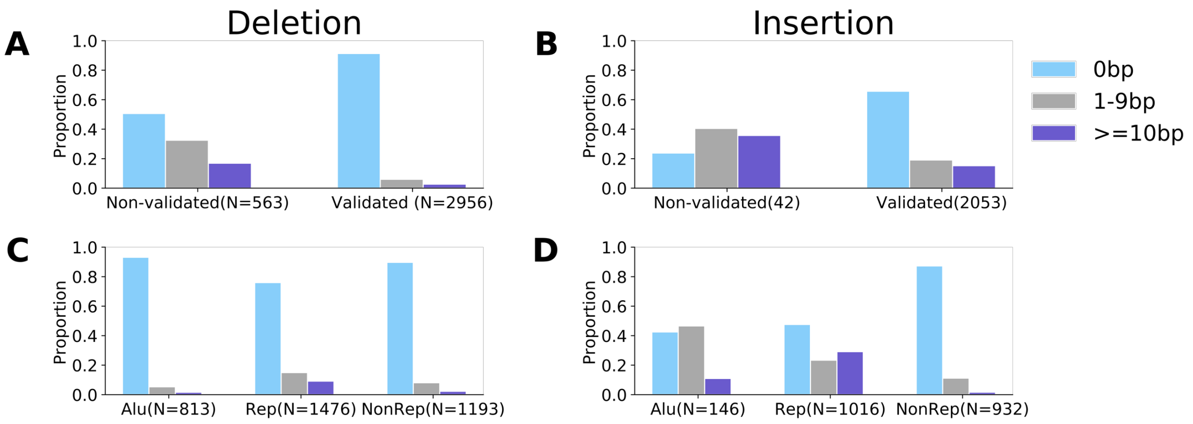


**Figure S14.** SV call size differences between NA12878 and NA24385. A and C for deletions; B and D for insertions. Non-validated: SVs are not validated in either of the two samples; Validated: SVs are validated in either of the two samples. Alu: SV matches Alu element consensus. Rep: repeats annotated by RepeatMasker. NonRep: SVs from non-repeats regions. The SVs were validated by PacBio reads and aligning to Ape genomes.
